# Muscle control of an extra robotic digit

**DOI:** 10.1101/2025.06.18.658087

**Authors:** Julien Russ, Lucy Dowdall, Francesco Cenciarelli, Kitty Goodridge, Mabel Ziman, Hristo Dimitrov, Dani Clode, Tamar R. Makin

## Abstract

Controlling an extra robotic finger requires the brain to adapt existing motor signals. While most current strategies exploit physical movement, there is growing interest in harnessing muscle activity directly via surface electromyography (EMG) as a more seamless interface. We systematically compared muscle- (EMG) and movement-based (force sensor) control of a Third Thumb. Using identical instructions and a counterbalanced within-participants design, we assessed initial skill, learning, and cognitive load across a variety of tasks, enabling a blinded comparison across control modalities. Both control modalities afforded successful Third Thumb control and learning, although force control consistently delivered better performance. Despite execution differences, learning rates and cognitive loads were comparable, with a similar evoked sense of agency. Signal analyses showed performance was predicted by real-time force sensor parameters but not by EMG, reflecting distinct control dynamics. Nonetheless, EMG training led to greater skill transfer to force control, suggesting it may better support generalisable learning. These findings challenge the assumption that proximity to neural signals ensures better control. Although EMG underperformed in execution, it showed unique advantages, including enhanced generalisation and access to richer signals, highlighting the need for improved real-time decoding to fully exploit its potential.

## INTRODUCTION

Recent advances in biotechnology, robotics and artificial intelligence have led to the rise of human augmentation technologies, aiming to enhance and expand human physical capabilities beyond their natural limitations (Prattichizzo et al., 2021; Eden et al., 2022). These devices take many forms, from artificial limbs or other end-effectors controlled by the biological body (Dominijanni et al., 2021) to technologies such as exoskeletons allowing enhanced abilities (Mooney et al., 2016; Kim et al., 2019). Beyond the motor domain, augmentation technologies may provide the user with cognitive enhancements such as smart glasses, providing tailored real-time data to a given context (Follmann et al., 2019), or neurostimulation devices (Waight et al., 2023). Augmentation technologies are not set in stone, their potential is vast with wide-ranging applications. This versatility implies that augmentation devices can be tailored to meet the needs of diverse user groups.

Supernumerary robotic fingers (SRFs) are advanced wearable devices that augment the human hand with additional robotic digits, enabling enhanced dexterity and functionality (Wu et al., 2014). Unlike prosthetics designed to replace lost biological function, SRFs coexist with the biological hand, therefore moving beyond replacement towards the enhancement of motor functions. While SRFs expand our physical capabilities, this extension introduces a bottleneck: the need for external control mechanisms, such as wearable sensors or neural interfaces, to operate the additional fingers without interfering with existing biological function (Prattichizzo et al., 2021; Dominijanni et al., 2021). A way around this key challenge is to exploit existing motor redundancies (Makin et al., 2022).

Motor output is shaped by multiple stages of the nervous system, from high-level neural commands to muscle activation. It is therefore possible, in theory, to tap into the motor system at all stages of the sensorimotor hierarchy to produce additional control signals, while maintaining existing motor functions. Non-invasive approaches like electroencephalogram (EEG) and electromyography (EMG) can capture neural and muscular activity that can be then repurposed to control an augmentation device (Verdel et al., 2024). While there are multiple methods to discern muscle activity, a simple surface electrode can yield a clear signal with high signal-to-noise ratio (Arozi et al., 2020), making EMG a popular choice for various controllable augmentation devices and prosthetics (Hahne et al., 2018; Hussain et al., 2016; Meraz et al., 2017; Tavakoli et al., 2017; Yang et al., 2014), as well as non-motor augmentation technologies (Trigili et al., 2019; Carvalho et al., 2023; Kiguchi et al., 2012), and other interfaces for gesture recognition or teleoperation (CTRL-labs at Reality Labs et al., 2024; Jie et al., 2021). However, to benefit from EMG for motor augmentation control, users must learn to reappropriate muscle signals intended for motor control of a specific body part to instead control a new body part (Dominijanni et al., 2021; Bräcklein et al., 2022; Formento et al., 2021). This raises the empirical question of whether users are better able to adapt their motor plan to best support EMG control, relative to other control sensors.

We wanted to test the accessibility of EMG control for a SRF with an emphasis on initial skill, skill learning, and intuitive control. As a model for studying the accessibility and precision of an EMG interface, we used the robotic Third Thumb (Figure 1), created by Dani Clode Design (Kieliba et al., 2021). The Third Thumb is a three-dimensional (3D) printed robotic thumb, designed to extend and enhance the motor abilities of an already fully functional hand (Figure 1). Originally, the Third Thumb was designed to be controlled by movement of the wearer’s toes via force sensors (FSs), providing proportional control across two degrees of freedom. With the toe-controlled interface, we previously demonstrated that individuals could perform basic motor functions almost instantly (Clode et al., 2024), allowing users to achieve skilled hand-device coordination and collaboration over a short time scale (Amoruso et al., 2022).

**Figure 1.**
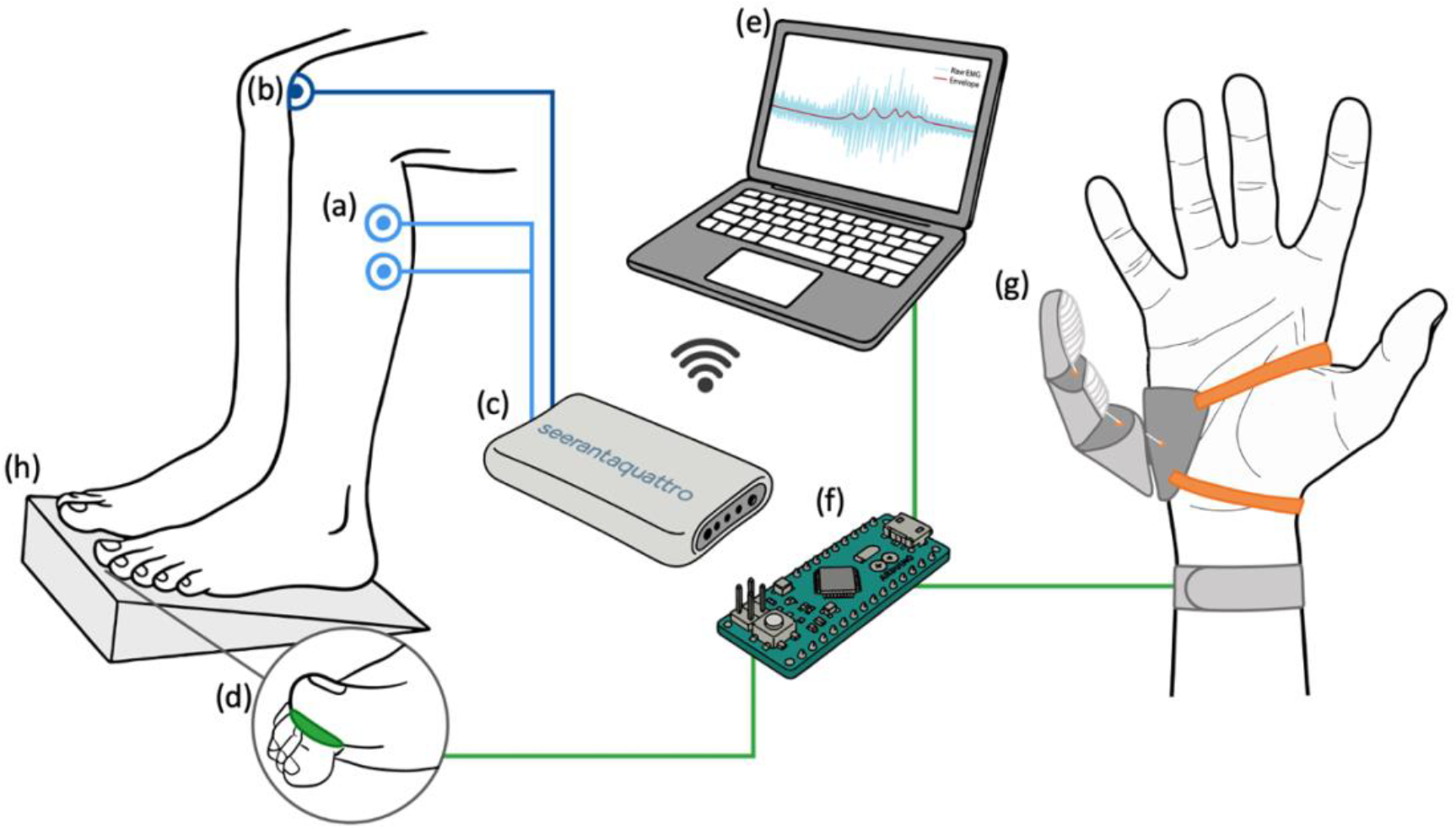
Study technical setup. (a) Bipolar electrodes placed on the gastrocnemius muscle of each leg to record the EMG signal. (b) Reference electrode to establish baseline. (c) Wireless amplifier to amplify and transmit the EMG data to the computer. (d) Force Sensors (FS) used to record pressure applied by the big toes. (e) Computer to convert control signals to Third Thumb motor commands. (f) Arduino receiving the commands from the computer and actuating the servos of Third Thumb. (g) The Third Thumb (Dani Clode Design). (h) Foot stand to constrain movements and co-contraction.

In the current study we replaced the traditional force control of the Third Thumb with EMG proportional control, also initiated by proportional toe movements. The EMG control was then put to the test by providing participants with identical instructions and experimental design, while they controlled the device using each of the two control methods in turn (see Figure 1 for technical setup). This counter-balanced design allowed us to directly compare performance between muscle and movement control across a variety of motor tasks. We hypothesized that EMG control will be more intuitive because it affords acting with the body, rather than acting through a sensor, i.e. an EMG interface is closer to the nervous system than the traditional toe sensors. We therefore predicted that EMG control will produce greater learning gains following brief training, as well as lower cognitive load during EMG motor control, and perhaps evoke a stronger sense of embodiment.

## RESULTS

### Behavioural Results

To compare muscle- and movement-based control, we used a within-subjects counterbalanced design. The experimental design is elaborated in Figure 2A. Participants completed a test-train-test sequence with one control method, then repeated it with the other modality. Training tasks targeted different motor skills–individuated Thumb use, Thumb–finger collaboration, or Thumb–finger coordination (Figure 2B). A proportional control task assessed baseline performance and post-training gains.

**Figure 2.**
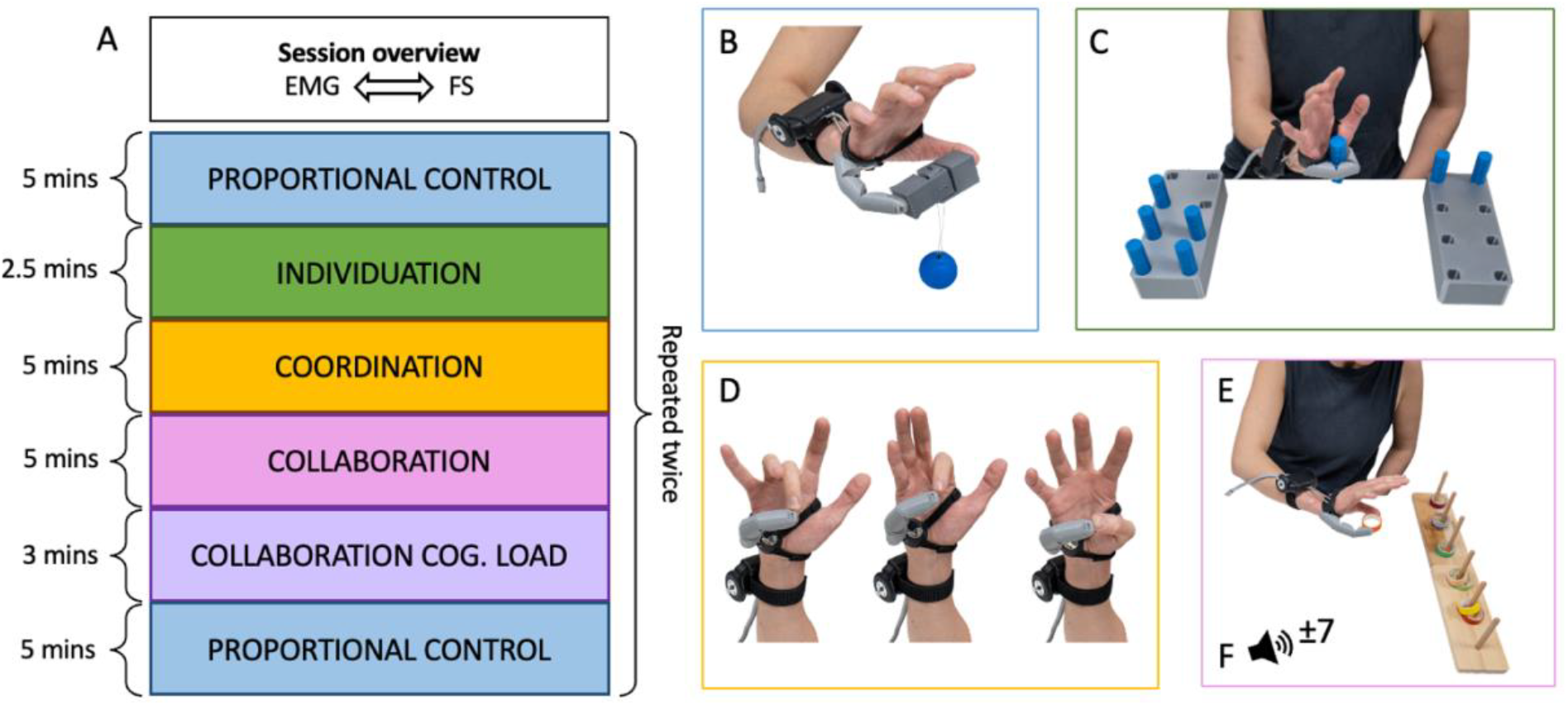
Study design. (A) Timeline for each task battery. Participants perform a series of training tasks, targeting different aspects of Third Thumb motor skill, with a Proportional Control task to assess training gains pre- and post-training. This battery was repeated twice for each participant (movement control and muscle control), with order of control methods counter-balanced across participants. (B) The Proportional Control task was repeated pre- and post-training. In this task, participants had to pick-up a rectangular ‘fragile egg’ of unstable mass (blue sphere) and transport it to a target location using the Third Thumb and biological thumb in collaboration. If too much pressure was applied, the egg would break. (C) In the Individuation training task, participants used the Third Thumb to transport pegs from one brick to another, placing the pegs into a designated slot. (D) In the Coordination training task, participants used the Third Thumb to oppose the tips of their biological fingers. (E) In the Collaboration training task, participants used the Third Thumb in collaboration with the biological thumb to pick up rolls of tapes and stack them over wooden pegs. (F) In the dual task, the Collaboration was repeated while participants simultaneously performed an arithmetic task, which was cued by tones.

### Movement control yields better performance, with comparable cognitive load across modalities

We first conducted a baseline analysis to confirm that participants could effectively control the Third Thumb using both control modalities. Participants successfully completed the pre-training Proportional Control task with both controls, as demonstrated by a significantly positive success rate [EMG: t(23)=6.803, p<0.001; FS: t(23)=11.487, p<0.001], however, excess force trial rates were high, indicating poor proportional control [EMG: t(23)=8.756, p<0.001; FS: t(23)=6.236, p<0.001] (Figure 3A-B).

**Figure 3.**
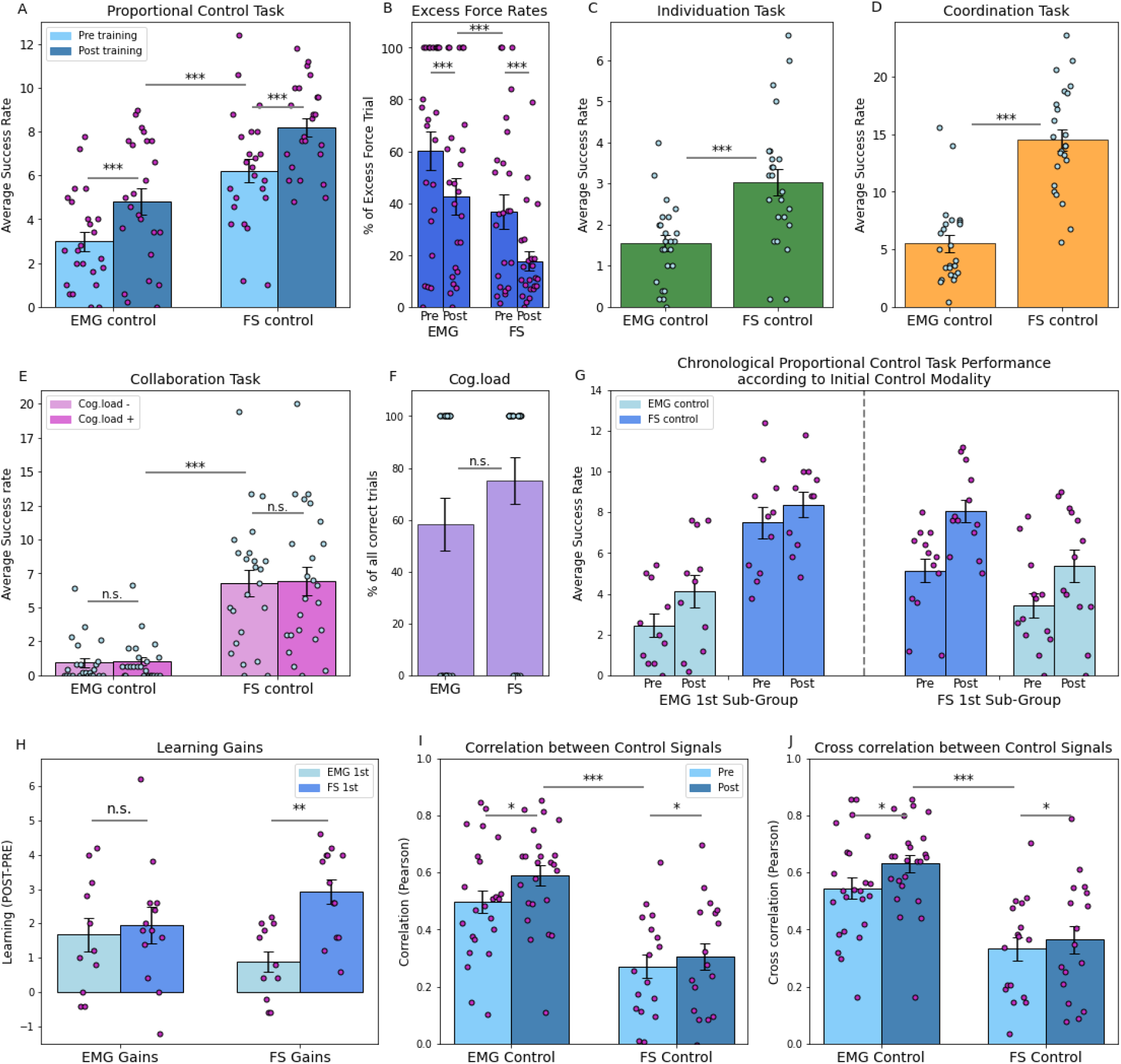
Behavioural task performance and relationship to control modality. (A) In the Proportional Control task, performed pre- and post-training for each control modality, FS control significantly outperformed EMG control, as measured using the average number of eggs moved (success rate). Performance significantly improved with training across both control modalities. (B) In the Proportional control task, the percentage of trials where excess force caused the egg to break was significantly reduced during FS control, with both control modalities showing improvement following training. (C) In the Individuation training task, the average number of pegs moved per minute (success rate) was significantly higher with the FS control than with the EMG control. (D) In the Coordination training task, the average number of finger oppositions per minute (success rate) was significantly higher in the FS control than the EMG control. (E) In the Collaboration training task, the average number of tapes moved per minute (success rate) was significantly higher for the FS control during both motor execution alone, and when there was an additional cognitive load. No deficits in performance were seen when a cognitive load was added for either control modality (BF_10_ = 0.295). (F) During the dual task, we extracted the percentage of blocks where the participant miscalculated the arithmetic. The average percentage of incorrect blocks was not significantly different across the control modalities (BF_10_ = 0.670). (G) In the proportional control task, learning gains were impacted by the order by which the two control modalities were performed, with participants starting with the EMG control (left) showing high pre-training performance using FS, whereas participants starting with FS control (right) show more modest EMG baseline performance relative to those starting training with EMG (left). (H) EMG training gains are similar regardless of starting control modality (BF_10_=0.383), but FS training gains are significantly higher for the FS-first sub-group. (I) Correlation between the EMG and FS control signals is significantly higher when participants are utilising EMG control, with a significant increase from pre to post testing sessions. (J) Cross correlation between the EMG and FS control signals showed similar results to correlation. Bar plots represent group means, error bars show standard error of the means, and single dots show individual participants. Asterisks denote significance as follows: * p<0.05, ** p<0.01, *** p< 0.001

Next, we compared performance between the EMG and FS control modalities across motor training tasks (Figure 3C-E). A significant advantage for the FS control was consistently observed throughout the training phase across all tasks (Individuation: t(23)=4.278, p<0.001; Coordination: t(23)=9.101, p<0.001; Collaboration: t(23)=6.624, p<0.001) (Figure 3C-E).

To investigate potential differences in attentional demands between control modalities, we examined the impact of additional cognitive load on motor performance. Introducing a concurrent arithmetic task during the Collaboration task did not significantly affect performance for either control modalities, compared to without the cognitive load (no interaction with control: [F(1,23)=0.024, p=0.877, BF_10_=0.413; post-hoc tests confirmed no difference for either control, EMG p = 0.749, BF_10_ = 0.284, FS p = 0.596, BF_10_ = 0.230)]. Counterintuitively, for both controls, participants exhibited a slight performance improvement when cognitive load was introduced (Figure 3E), likely reflecting gains from motor task repetition, as the cognitive load task was always presented after the Collaboration task. These findings suggest that neither control modality imposes substantial cognitive demands for producing Third Thumb movements. Additionally, arithmetic task performance did not significantly differ between the two control modalities (χ^2^(1)=0.229, p=0.633, BF_10_=0.562; Figure 3F), further supporting the notion that their cognitive demands are comparable, although the evidence remains anecdotal.

### Similar training gains across control modalities, despite superior movement control performance

To investigate learning effects across control modalities, we examined training gains between the pre- and post-training Proportional Control tasks. Here, too, we found clear evidence for a superior success rate for movement-based FS control (F(1,22)=41.206, p<0.001), which was apparent during both baseline (t(28.431)=5.905, p<0.001) and post-training (t(28.431)=6.069, p<0.001). Superior FS performance was also demonstrated when comparing proportion of excess force trials (F(1,22)=14.388, p<0.001). Regardless of the control modality, participants exhibited a significantly higher success rate following training (F(1,22)=65.385, p<0.001), with comparable training gains across both modalities (EMG: *M*=1.83; FS: *M*=1.99; t(23)=0.378, F(1,23) = 0.143, p=0.709, BF_10_ = 0.229). Correlations between pre- and post-training success rate were similar for both control methods (EMG: R=0.806, p<0.001; FS: R=0.805, p<0.001), indicating inter-individual variability in performance was stable across training. Notably, at the end of the brief training session, EMG control did not achieve similar performance to FS control, reinforcing the superiority of FS in task performance.

### Muscle control is a better tutor for Third Thumb motor skill

It is well established that repeated exposure to task requirements leads to performance gains (Makin et al., 2019). We would therefore expect participants to exhibit baseline gains when repeating the tests battery in the second part of the study, regardless of the control modality. This provides a unique opportunity to examine learning transfer effects–– whether movement-based training enhances muscle control and vice versa.

For each participant, we re-analysed movement- and muscle-control success rates at both pre- and post-training for the Proportional Control Task, incorporating the order in which the control modalities were introduced as a between-subject factor (Figure 3G). While the two-way interaction was not significant (Control X Pre/Post: F(1,22) = 0.052, p = 0.822, BF_10_ = 0.443), a significant three-way interaction was observed (F(1,22)=4.521, p=0.045), indicating that control impacts pre-to-post learning gains, depending on the control modality used initially. This suggests that the transfer of learning between EMG and FS is asymmetrical––one control modality serves as a more effective tutor than the other. This is a potentially important observation, as it hints that control-invariant task repetition alone cannot fully explain improvements across the training batteries.

Figure 3G shows performance differences between the two sub-groups of participants starting the experiment with muscle control (left; EMG training first) versus those starting training with movement control (right; FS training first). People starting the session with EMG training showed higher performance when starting the FS session (FS pre; left side), compared to those starting with FS from scratch (FS pre; right side of Figure 3G) (t(22)=2.374, p=0.027). Whereas, participants who trained with FS first and then switched to EMG (EMG-pre; right) exhibited more comparable pre-training performance with EMG to those starting with EMG training first (EMG-pre; left) (t(22)=1.115, p=0.277, BF_10_=0.586). This suggests greater generalisation of learning from EMG to FS control. However, the FS learning gains from pre-to post-training is markedly lower for participants who started with EMG (t(22)=4.083, p<0.001; Figure 3H). This observation is likely facilitated by ceiling effects, limiting FS control learning gains from pre-to post-training as performance had already reached a high success rate following the initial EMG training (Figure 3G-H).

To further explore why muscle control is a better tutor, we correlated control signals for the main degree of freedom (flexion/extension) over time across modalities. Using both correlations and cross-correlations (Figures 3I–J), we find that EMG and FS signals are highly correlated while participants are using the EMG output to control the Third Thumb, with significantly higher correlations than during FS control (F(1,16)=29.140, p<0.001). Therefore, even when using the EMG control signals, participants are expressing the FS-related toe movement, hence contributing to relevant task-learning. Interestingly, this association between control modalities further grows with training, as indicated by a significant increase in correlation from pre to post-training (F(1,16)=5.541, p=0.032), with no interaction effect based on control order. Performing the same analysis with cross-correlation (Supplementary Figure S1), to account for potential delays between the two signals, we found similar effects (control modality effect: F(1,16)=31.589, p<0.001; learning effect: F(1,16)=4.962, p=0.041). These findings provide a mechanistic insight into the high generalisation capacity of EMG training.

### FS control signal parameters can explain behavioural results

Because both EMG and force signals serve as interfaces between the user and the robotic device, their structure and dynamics may influence control quality. To better understand sources of performance variability, we analysed the control signals themselves. We extracted a set of descriptive parameters from both the EMG and force signals, such as signal amplitude, smoothness, and complexity, reflecting different aspects of signal behaviour, from stability to temporal complexity. These parameters, detailed in the Methods section and Supplementary Figure S2, were used to investigate the underlying reasons for the movement-based control advantage, as well as the bidirectional transfer of learning between EMG and FS control.

### Force signal is more directly linked to task performance

We first assessed whether differences in the real-time FS control parameters could predict inter-individual differences in movement control task performance. We focused on post-training performance in the Proportional Control task, as a measure of users’ control proficiency after initial learning. To determine whether signal parameters or participants’ skill better explained performance (or a combination of both), five of the eight computed parameters–mean, standard deviation, fractal dimension, slope change, and second derivative variance–along with behavioural performance during the EMG phase (with control order as a covariate), were included in a backward stepwise regression (see Supplementary Table S3 for full regression output). The model at step 3 demonstrated the best fit to the movement control behavioural data, explaining 58% of the inter-individual variance (p=0.004), with significant contributions from several FS parameters (fractal dimension (p=0.014), second derivative variance (p=0.003), mean (p=0.070)), as well as EMG performance (p=0.009) and control order (p=0.005). As both control and performance parameters were significant predictors, these results indicate that both signal characteristics and individual skill contribute to performance under movement-based control.

We repeated the same analysis for EMG performance using the five EMG control parameters (mean, standard deviation, fractal dimension, slope change and second derivative variance) alongside behavioural performance during the force control phase (with control order as a covariate). The model did not significantly converge, with the best model at step 6 explaining 10% of the variance (see Supplementary Table S4 for full regression output). None of the EMG parameters were significant predictors of muscle-based control, while behavioural performance under force control being the strongest predictor, trending towards significance (p=0.085). Therefore, EMG signal control characteristics alone, or with the additional behavioural parameters, do not sufficiently account for performance under muscle-based control.

These findings suggest that, relative to EMG, FS control is more directly linked to behaviour. As the extracted control parameters do not provide a sufficient or direct mapping to task execution, this likely contributes to the poorer performance with EMG control.

### Raw EMG signal contains performance-relevant parameters

For completion, we investigated whether raw EMG signals (the initial unfiltered control signal produced by participants) could better explain muscle and force control performance than the filtered control signal.

Repeating the analysis as above for EMG performance, using the same five parameters, but computed from the raw EMG time series (fractal dimension, mean, standard deviation, zero crossing, and slope change) alongside force control performance (with control order as a covariate) yielded a strong model at step 2, explaining 41% of the variance (p=0.030). Notably, force control performance (p=0.009), EMG parameters (slope change (p=0.007), zero crossing (p=0.011)), as well as control order (p=0.015) all showed significant contributions. The model also contained non-significant contributions from the mean (p=0.191) and standard deviation (p=0.088) (see Supplementary Table S5 for full regression output). While the resulting model is a combined contribution from behavioural information and control parameters, this finding still suggests that raw EMG has multiple potential factors that strongly predict behaviour but are being filtered out or not entirely captured when converting the raw EMG to the control signal. Hence, processing the raw EMG signal to the control signal resulted in a loss of information, suggesting the aspects of the signal utilized for control are suboptimal.

Finally, when predicting movement control performance using raw EMG parameters, we obtained a strong model fit (38% variance explained, p=0.017). The final and best model (step 5) contained EMG slope change (p=0.015), EMG performance (p=0.008), and control order (p=0.022), where all three were significant predictors (see Supplementary Table S6 for full regression output). This result reveals that during the FS task, participants meaningfully engage the muscles which were also exploited for EMG control.

### Greater shared information between the two degrees of freedom during EMG control

We next focused on coordination between the Third Thumb’s two controllable degrees of freedom (DOFs) during the Proportional Control task. We used distance metrics–mutual information and dynamic time warping (DTW)–to assess whether one control modality facilitated more coordinated co-usage. We found significant evidence that mutual information was higher during EMG control (F(1,18)=7.186, p=0.015) (Figure 4A). This effect suggests greater shared information between the two DOFs during EMG control. There was no significant difference in mutual information from pre to post-training (F(1,18)=1.893, p=0.186, BF_10_=0.122). Conversely, DTW distance was significantly lower for force control compared to EMG (main effect of control modality: F(1,18)=5.952, p=0.025; Figure 4B), indicating more synchronized and consistent movements during force control. While mutual information did not significantly differ between pre- and post-training across controls, there was a trend towards increased DTW distance from pre to post (F(1,18)=3.476, p=0.079). This suggests that participants increasingly adopted a sequential activation strategy for the two DOFs following training. These findings highlight key differences between the two control methods: EMG appears to facilitate simultaneous use of the DoFs (higher mutual information), while FS enables more precise timing (lower DTW). However, across the experiment, the emerging strategy trended towards sequential, and presumably more accurate, movements.

**Figure 4.**
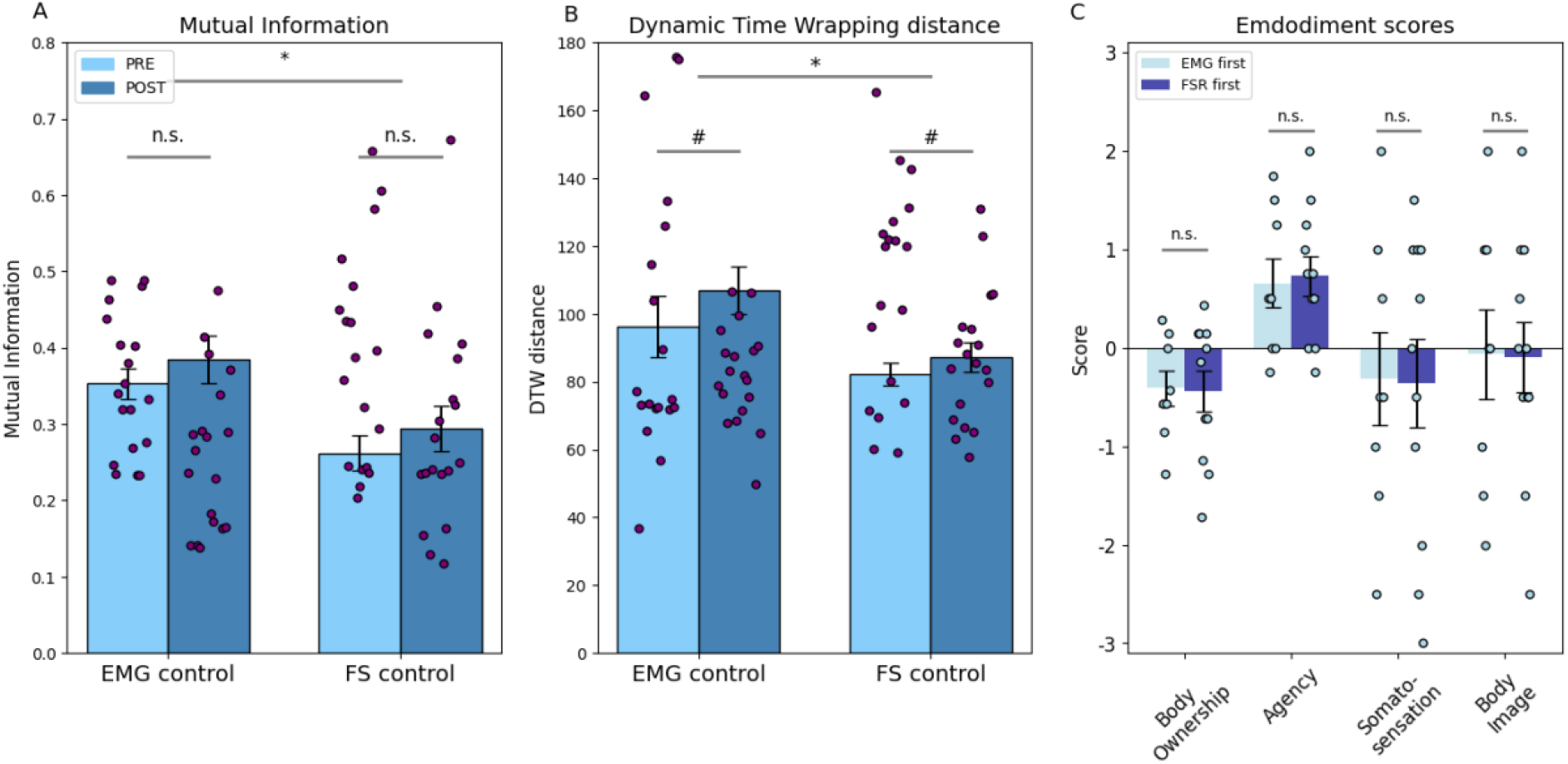
Coordination analysis between the two degree of freedom (left and right leg) and embodiment scores. (A) For the Proportional Control task, mutual information is significantly higher during EMG control. (B) For the Proportional Control Task, dynamic time wrapping (DTW) distance is significantly higher for EMG control, additionally there is a increasing trend from pre-to post-training. (C) Embodiment scores across the four explored categories, show similar ratings across control modalities (EMG or FS first). Positive ratings, indicating perceived integration with the body, were found for sense of agency, whereas ownership was negative, indicating the Third Thumb was perceived as an external robotic device. Bar plots represent group means, error bars show standard error of the means, and single dots show individual participants. * denotes p<.01, # denotes p<.08

### Both control modalities induced a sense of agency, but not body ownership

Lastly, we examined participants’ subjective experience of embodiment with the Third Thumb, using four standard dimensions: agency, body ownership, somatosensation, and body image (Figure 4C). To avoid response contamination, the embodiment questionnaire was administered after the first control session. This design allowed for a between-subjects comparison of control modality (muscle versus movement). Since no significant differences were found between control sub-groups across any of the embodiment measures (agency: t(17)=0.214, p=0.833, BF_10_=0.414; body ownership: t(17)=0.103, p=0.919, BF_10_=0.409; somatosensation: t(17)=0.073, p=0.943, BF_10_=0.408; body image: t(17)=0.047, p=0.963, BF_10_=0.408), we report combined results below; individual sub-group statistics are reported in the Supplementary Materials.

Participants reported significantly positive agency scores (t(18)=4.373, p<0.001), whereas body ownership scores were consistently negative (t(18)=2.991, p=0.008), indicating participants perceived the robotic finger predominantly as an external object, but nevertheless perceived that they could control it. Somatosensation and body image scores were neutral (t(18)=1.013, p=0.325, BF_10_=0.373; t(18)=0.273, p=0.788, BF_10_=0.246, respectively), reflecting neither positive nor negative interpretation of related statements.

Overall, these findings highlight agency as the only positively rated dimension, with no substantial differences between the EMG and FS control methods.

## DISCUSSION

Our findings portray the distinct strengths and limitations of movement- (FS) and muscle-based (EMG) control, positioning movement control as a more immediately effective interface while revealing muscle control’s potential role in augmenting motor learning.

Our findings demonstrate that, irrespective of the control modality, participants were able to successfully perform the motor tasks and exhibited a clear trajectory of improvement over the course of each session. This suggests that both modalities enable users to acquire and refine control over the augmentation device. Despite overall task proficiency, a consistent advantage of FS over EMG was observed across all assessed motor skills. This superiority may be attributed to the more direct and stable signal acquisition of FS, reducing variability in control and allowing for finer adjustments of movement execution. Furthermore, predictive modelling revealed that FS control parameters reliably explained performance outcomes, whereas EMG parameters did not. Instead, EMG performance was best predicted by behavioural performance while participants were using movement control, reinforcing the notion that muscle control introduces additional complexity. The robustness of movement control highlights the effectiveness of pressure-based control in motor tasks across the range of motor tasks tested here. Nevertheless, muscle control serves as a more effective tutor for generalisation of motor control, facilitating greater improvements in subsequent FS control.

Despite consistently superior performance with force control, we found no significant differences in cognitive load or overall embodiment between the two control modalities. The introduction of an additional arithmetic task did not result in performance deficits. Across the embodiment dimensions, participants’ perception of the Third Thumb was comparable across tasks. Together these findings highlight that despite its lower motor performance, muscle control does not inherently decrease the cognitive resilience or the sense of control. However, the lack of a stronger relationship between agency and control modality could be attributed to participants’ limited prior experience with either modality: without prior experience, any form of control might be sufficient to produce a positive subjective sense of agency. Regardless, given that previous studies reported increased sense of body ownership following longitudinal training to control the Third Thumb (Kieliba et al., 2021), prolonged experience and refinement of motor strategies may elicit a more distinct sense of embodiment in one of the two modalities.

Why was movement control superior to muscle control? Several key factors contribute to the superior performance of FS over EMG. Fundamentally, pressing the toes to apply force– required for FS activation–is a more intuitive and straightforward action than selectively contracting a given muscle for EMG control. Motor learning is typically feedback-driven, as we naturally adapt movements based on interaction with the physical world (Scott et al., 2015). The articulation of a movement against a surface provides richer and therefore more distinct haptic feedback to support motor control and learning (Amoruso et al., 2022). Furthermore, force sensors directly measure force application, providing a stable and proportional control signal that better aligns with the desired augmentation control paradigm of this study. In this context, the high correlation between control signals during EMG control supports that, while there are multiple ways to ‘solve’ FS control, there are fewer routes to learning to ‘solve’ the EMG control challenge.

In contrast, muscles function in synergies, making it difficult and unintuitive to isolate and control a single muscle for fine motor tasks (d’Avella et al., 2003). In addition, muscle activation does not inherently follow the external feedback model. As such, unintended leg movements–despite being constrained with a foot stand–could affect EMG signals more than FS signals. Then, the nature of motor unit recruitment for muscle activation further complicates EMG control. Unlike movement, which is continuous and proportional, muscle engagement is discrete and occurs in step-like increments. Motor units fire in accordance with recruitment patterns (Farina et al., 2002), leading to inherent variability even under constant contraction. This variability is exacerbated by the time-varying nature of EMG signals (Stegeman et al., 2000), which are not inherently suited for proportional amplitude recording. Even when attempting to maintain a steady muscle contraction, EMG signals can fluctuate due to physiological and electrode placement factors, introducing unintended variations in control (Wong et al., 2006). These considerations suggest that EMG control involves more complex and dynamic motor strategies, making it inherently more challenging to master.

Another crucial factor is signal processing. While FS signals are averaged over time and directly mapped to an output angle with minimal transformation, EMG control relies on a signal feature–amplitude–which is smoothed (from the raw EMG) and time-averaged, reducing the directness of control. Using amplitude from a single EMG channel, although intended to closely mimic the straightforward approach of FS control, may be also suboptimal due to limitation in signal filtering. This difference in signal reliability is reflected in our predictive modelling results, where FS performance could be explained using mean force values, i.e. the average amount of pressure applied was enough to explain participants’ motor performance; while EMG performance was best predicted by transient features such as slope change and zero crossings– indicative of fluctuating muscle activations rather than steady control. Nevertheless, it is worth pointing that the raw EMG signal could be a rich source of task-relevant information, as demonstrated when predicting both FS and EMG performance using the raw EMG signal for off-line analysis.

Interestingly, it is because EMG is a more challenging interface to adopt when first learning to control a motor augmentation device, that it might also be a more efficient tutor for generalised motor learning across modalities. Although counterintuitive, this effect aligns with existing literature suggesting that more challenging training conditions foster greater generalisation across motor contexts (Schone et al., 2024). Additionally, studies in cognitive neuroscience emphasise that learning from more variable input conditions, while slower initially, results in better and more robust generalisation (Raviv et al. 2022). Thus, despite lower immediate performance, EMG training might enhance the cognitive and motor strategies necessary for effective generalisation. An alternative interpretation to this greater generalisation of learning is rooted in the greater correlation observed between EMG and FS signals during muscle control. When participants are operating the EMG control, they are producing task-relevant movements with their toes. This could be due to the challenges of intuitive muscle control discussed above, or due to our explicit task instructions, which emphasised movement. Either way, by exercising toe movements during EMG control, participants are implicitly training a movement strategy that will benefit their performance while switching to the new control modality. Either way, this generalisation of motor learning raises the possibility that EMG training is advantageous for preparing users for varied motor control tasks and scenarios.

Several limitations in our study design must be acknowledged. Our training sessions were relatively short and learning gains were comparable between muscle and force control. With more extensive training, EMG could potentially match or even surpass FS performance, particularly as participants already showed signs of approaching ceiling levels with FS control. This aligns with evidence suggesting that while varied inputs such as EMG may result in slower initial learning, they do not inherently limit ultimate performance (Raviv et al., 2022). Additionally, our device and training paradigms were specifically tailored to FS control, potentially biasing performance outcomes in its favour. Furthermore, the hardware approach of using a single EMG channel per degree of freedom to mimic FS control likely limited EMG effectiveness; future approaches could greatly enhance EMG-based augmentation technologies by employing high-density EMG arrays, providing more reliable amplitude data. Finally, software improvements–such as integrating more dynamic parameters like slope changes or zero crossings or utilizing minimally processed EMG signals–may further boost EMG’s practicality and efficacy in real-world augmentation scenarios.

## Conclusion

Our findings suggest that, while EMG-based muscle control provides access to neural signals closer to the source of movement generation, its inherent variability and complex processing pipeline make it less effective for precise proportional control compared to movement-based force sensing. Although our initial hypothesis posited that control would improve as the interface approached the nervous system, our results challenge this assumption, demonstrating that a more distal, movement-based control can yield superior early performance. Nevertheless, our results clearly demonstrate that EMG, although currently not superior to FS control, remains a viable modality for augmentative motor control, with participants successfully operating the Third Thumb using only a single bipolar electrode per degree of freedom. Addressing EMG preprocessing limitations through optimised hardware and advanced signal-processing techniques could significantly enhance EMG’s practical applicability in augmentation technologies. As such, we see EMG control as a highly promising future technique for motor augmentation control.

## METHODS

### Participants

Twenty-four healthy volunteers were recruited to participate in the study via the MRC Cognition and Brain Sciences and Department of Psychology SONA participant pools at the University of Cambridge, as well as via word of mouth. Participants were randomly allocated to the EMG control first (6 females, 5 males; Age *M*=21.9 (*SD*=3.5)) or force control first (6 females, 7 males; Age *M*=25.5 (*SD*=8.6)) sub-groups. All participants were right-handed, and did not have any known motor disorders. Ethical approval was granted by the Cambridge Psychology Research Ethics Committee (PREC: 2022.068). All participants were provided with an information sheet and gave their written consent before participating in the study. Eight more participants had their data collected but were excluded from analysis due to technical error or incomplete dataset. Five participants did not complete the embodiment questionnaires.

### Third Thumb

The Third Thumb is a 3D printed supernumerary robotic finger (Dani Clode Design), designed to extend and enhance the abilities of a fully functional hand (Kieliba et al., 2021). The Thumb is worn over the ulnar side of the right palm, opposite the user’s natural thumb (Figure 1). It is actuated by two servo motors, allowing proportional control of two independent degrees of freedom (DOF), flexion/extension and adduction/abduction. The motors are mounted on a wrist strap and in our setup powered by a wall plug (see full description in Clode et al., 2024; see Kieliba et al., 2021 for description of wireless setup).

### Technical Setup

1. **Force control Thumb** In the original Third Thumb design, control signals are generated from the user’s toes using force sensing resistors, which vary their electrical resistance based on applied pressure. A wired communication protocol sends signals from the controlling sensors to the motors to actuate the Thumb. Signals from the right toe control the Thumb’s flexion (pulling the Thumb across the hand) while signals from the left toe control adduction (pulling up towards the fingers). Thumb movements are fully proportional to the control signal, allowing the user to vary its position by modulating the force applied to the sensors. The study’s technical setup is displayed in Figure 1. For the FS setup, a sensor was fixed to the underside of each of the user’s big toes. Prior to starting the experiment, a calibration process was performed to ensure the full range of Third Thumb motion was accessible to the participant. Participants were asked to rest their feet on a footrest before pressing hard with both of their toes for approximately 10 seconds. For each sensor, 60% of the maximum sensor value was then used as the ceiling. This prevented potential fatigue that could reduce the accessibility over the full dynamic range of the Thumb (all possible positions). To extract commands from the FSs, values were sampled at a 2000Hz sampling rate, normalized according to the calibration range, and then appended to a buffer of 64 sample points (around 300ms) that acted as a moving average to ensure smooth movements. Each DoF had its own buffer. The buffers average was updated every 20ms and mapped to the dynamic range of each DoF (160° for flexion and 90° for abduction).
2. **Muscle control Thumb** For the EMG setup, two bipolar electrodes were placed on the gastrocnemius muscles (medial head) of each leg (Figure 1). This muscle was chosen as it contracts when applying downwards pressure with the toes, enabling similar motor control to the movement control paradigm. An additional dry reference electrode was placed on the patella bone (knee). To reduce participants’ co-contraction of agonists and antagonist muscles during EMG control, a wooden support was designed to keep the feet at an angle of 20°, also constraining variability in EMG control styles between participants. Similarly to the force control, a calibration process also occurred at the beginning of the session to determine the EMG control range for each participant. During calibration, participants were asked to stay still and limit any contraction in their lower limbs (rest), before being prompted to contract their big toes against the foot support as hard as possible for approximately 10 seconds. 60% of the maximum filtered amplitude was taken for each foot respectively to determine the dynamic range ceiling for each DoF. The EMG sampling rate was 2000Hz, with all the electrodes wired to an amplifier (Sessantoquattro+, OTBioelectronica) that sent the raw data via a WiFi connection every 100ms to a computer (Dell 5820, Windows 10 v20H2). The raw EMG signal was first processed to extract its envelope by band-pass filtering the signal between 20Hz and 500Hz, band-stop filtering at 50Hz (to remove power line noise), followed by rectification and low-pass filtering at 2Hz. Each DOF of the third Thumb was controlled by a moving average (400ms) of the EMG envelope of the respective leg. The average of the buffer was mapped to an angle for the Thumb actuation based on the minimum (resting level) and maximum (60% maximum contraction) values of the calibration range.

### Experimental design

To assess performance differences between the two control methods (muscle and movement), a within-subjects counterbalanced study was designed. Participants underwent a test-train-test task sequence using the first control for approximately 30 minutes, and then switched to the second control method and repeated the test battery (Figure 2A). Both control modalities were setup at the beginning of the experiment, allowing us to blind the participant to which control they were using for a given task. Their only instruction was that pressing on the toes will actuate the Thumb proportionally. Following the first control method, participants completed an embodiment questionnaire.

The training tasks developed different motor skills, requiring either individuated use of the Third Thumb, collaboration between the Thumb and the biological fingers, or coordination of movements between the Thumb and the fingers. Before and after these training tasks, a ‘testing’ task focused on proportional motor control was used to determine baseline motor performance and then training gains.

### Testing task

For the testing task we used a modified version of the fragile virtual egg task (Schone et al., 2024; inspired by Clemente et al., 2016) which is designed to probe proportional control in a dexterous task (Figure 2B). The Thumb was used in collaboration with the biological thumb to pick up and transport an object (the ‘egg’) with unknown and unstable mass. The egg was cuboid-shaped, 8×3×3cm in dimension with a hanging spherical weight (5g), and weighs a total of 36 grams. It is called an ‘egg’ due to its mechanical property to collapse under too much load (collapsing at 2.7 newtons, ‘breaking’). Participants were asked to transport the egg (placed to their right) into a basket on their left (25×19×14cm) as many times as they could during one block repetition (lasting one minute). Participants were instructed to transport the object without dropping it or ‘breaking’ it. This task was repeated over five blocks, both before and after a twenty-minute training paradigm.

Outcome measures focus on early and late performance (pre/post-training), including average successes (average number of successful trials per block) and errors (average number of trials per block involving excessive force resulting in a ‘broken’ egg). We also measured learning gains on these outcomes by comparing performance before and after training.

### Training tasks

The training period comprised of three tasks, each focused on different motor skills when operating the Third Thumb. The tasks were performed in a set order:

1. **The Individuation task** (Figure 2C) was focused on motor control of the Third Thumb without support of the hand. The task consisted of eight 3D-printed cylindrical pegs (2.3×8.3cm, 17 grams, placed on a holder to the participants’ right) and another eight-slot peg brick holders (23.5×10×5cm) placed on their left. Participants were asked to move each peg to its corresponding slot on the empty brick and were given 30 seconds per block to move as many pegs as possible. In the event of a peg falling during transport, it was placed back to its starting position on the right. If the participant was able to slot all eight pegs before the block ended, additional pegs and a new set of brick holders were provided to continue the block. This task was performed for five block repetitions. The outcome measure was the average number of pegs successfully slotted in the brick holder across blocks.
2. **The Coordination task** (Figure 2D) required a coordinated finger opposition between the Third Thumb and a biological finger, with the movements resulting in tip-to-tip contact. Starting with the biological thumb and moving sequentially around the hand to the pinky, participants had to wait for the experimenter to confirm each trial as successful, before moving to the next finger. Once the participant reached the pinky finger, they started a new sequence beginning from the thumb. This task was performed for five blocks of one minute each. The outcome measure was the average number of finger oppositions across the five blocks.
3. **The Collaboration task** (Figure 2E) involved continuous collaborative engagement of the Third Thumb and biological hand. The task consisted of rolls of tape (3.8×1.5cm, 4 grams), placed to the participants right, and a series of six wooden poles on their left (7.5cm apart). Participants had to grab a roll of tape using the Third Thumb in collaboration with the tip of the biological thumb, and stack it onto a pole. Participants were asked to move as many tapes as they could in a minute, and the block was repeated five times. The outcome measure was the average number of tapes moved across the five blocks.
4. **The cognitive load task** (Figure 2F) consisted of two simultaneous components. The Collaboration task was repeated, and simultaneously with motor performance, participants were instructed to complete an arithmetic working memory task, designed to probe cognitive load during artificial limb motor control (Kieliba et al., 2021; Witteveen et al., 2012; Guthrie et al., 2022). At the start of each block, a random number, between 100 and 300, was given to the participants. Participants were then presented with a series of low (100 Hz) and high (800 Hz) pitch tones randomly occurring every 5-6 seconds. They were required to verbally respond to each tone by adding seven following a high-pitch and subtracting seven following a low-pitch tone. Participants were instructed to prioritize correct arithmetic task performance, with the motor collaboration task being secondary in importance. This dual task was repeated for three blocks, each lasting one minute. There were two outcome measures, the average number of tapes moved across the three blocks and the number of blocks where the participant did not answer the arithmetic task correctly.

### FS and EMG Parameters extraction

To deepen our insight into the control signal parameters that best predict successful motor control of the Third Thumb, a total of eight parameters were explored off-line, covering different aspects of the FS and EMG signals: The mean reflects the overall activity level of the muscle, indicating sustained contraction strength or effort. Standard deviation measures the signal’s dispersion, indicating the extent of variability in pressure/muscle activation levels. Variability refers to the changes in signal amplitude over time, reflecting the consistency of control/muscle contractions. Fractal dimension assesses the complexity of the control signal, providing insights into the intricate patterns of muscle activation. The first derivative variance captures the rate of change in the signal, highlighting the speed and smoothness of control signals. The second derivative variance measures the acceleration or deceleration in these changes, offering deeper insights into the dynamics of control and coordination. The slope change reflects the dynamics of signal variations and can indicate abrupt changes or events in control activity. Waveform length is the cumulative sum of the absolute differences between consecutive control signal sample points, often used to assess muscle fatigue or effort.

Five additional parameters were investigated for only the raw EMG signal: Frequency content involves the analysis of the signal’s spectral components, revealing the different frequencies at which muscles are active, which can be linked to different types of muscle fibre recruitment. The zero-crossing rate is the number of times the EMG signal crosses the zero-amplitude line within a time window (here one second), providing further insight into signal complexity in muscle activation patterns. The wavelet energy (complex Morlet wavelets were used for analysis) is the total energy of the EMG signal within specific frequency bands (here all frequencies present in the signal), obtained using a wavelet transform, highlighting localized energy patterns in time and frequency. Band energy is the same analysis targeted solely to the EMG frequency range (20 to 150Hz). Wavelet entropy is a measure of the disorder or randomness in the wavelet coefficients of the EMG signal, capturing signal complexity and irregularity, often used to differentiate between voluntary and involuntary muscle activity. The FS signal being strictly positive (as the rectified EMG signal) meant there was no need to look at zero crossings, and EMG specific parameters such as the wavelet energy or entropy that are not relevant for the force sensors signal.

When computing the FS/EMG parameters, the FS/EMG time series for each block of the Proportional Control task were extracted, all the parameters were then computed through a Python script. Parameters were averaged over block repetitions, resulting in one value per parameter for a given task. To minimise multiple comparisons, all candidate parameters were first correlated with each other. Similarity profiles were determined by how highly a parameter correlated with all the others (Supplementary Tables S1 and S2). Parameter with similar profiles were eliminated, isolating five unique signal characteristics that were used for statistical analysis (standard deviation, mean, zero-crossing rate, second derivative variance and fractal dimension.

### Coordination Parameters

To analyse coordination between the two DoFs, mutual information and dynamic time wrapping distance between the two DoFs were computed. **Mutual Information (MI)** is a method that captures both linear and nonlinear dependencies between the two input signals. If the mutual information is high, it suggests that there is a strong interaction or relationship between the two inputs, indicating synchronized coordination. **Dynamic Time Wrapping (DTW)** is a technique used to compare the similarity between two time series, even if they are not perfectly aligned in time. This can be useful if participants coordinate the two DoFs but with slight timing discrepancies (e.g. one DoF slightly lags behind the other). Extraction of the coordination parameters was done the same way as for the FS and EMG control signal parameters.

### Embodiment Questionnaire

To assess participants’ phenomenological experience with the Third Thumb, participants were asked to complete an embodiment questionnaire after the train-test battery with the first control. Participants were asked to rate their agreement with 20 statements (adapted from Kieliba et al., 2021; inspired by Longo et al., 2008) on a seven-point Likert-type scale ranging from −3 (strongly disagree) to +3 (strongly agree). Four statements were ‘catch’ statements, designed to confirm the participant was paying attention to their ratings. The other statements were clustered into four main categories, probing different aspects of embodiment, namely, body ownership, agency, body image, and somatosensation. For each participant, questionnaire scores were averaged within each embodiment category. Note that we did not repeat the questionnaire after the second control strategy, to avoid response contamination.

## Statistical Analysis

Data was processed using python (version 3.1713), edited in Visual Studio (VS) Code (version 1.87.2). All statistical analysis were performed in JASP (version 0.18.3) or using Python, edited in VSCode. All data was inspected for violations of the normality assumption by the Shapiro-Wilk test, and outliers were removed at a threshold of +/-3SD. An alpha level of .05 was used throughout, and any nonsignificant results of interest were followed-up with a corresponding Bayesian test (Cauchy prior width r = 0.707). Bayes factors were interpreted using BF<1/3 as sufficient evidence in support of the null hypothesis, BF>3 as sufficient evidence in support of the alternative hypothesis, and 1/3<BF<3 as inconclusive evidence (Wetzels et al., 2012).

When making pairwise comparisons of performance in the training tasks between the two control conditions, a student’s paired samples t-test was used. A repeated measures (RM) ANOVA was also used to assess performance in the dual task, with factors of control method and task version (collaboration or dual task). A chi-squared test was used to then further explore the proportion of trials in the dual task that had incorrect answers. A RM ANOVA was used for the pre-post testing task outcomes to compare training gains in each control condition, with a mixed ANOVA also used when considering control order. This was run separately for success rate and percentage of trials with excess force. Similarly, a RM ANOVA was used to compare correlation values between the control signals. Pearson’s correlation was used to correlate signal parameters with performance. Paired samples t-test were used to compare gains in the proportional control task with the two control modalities. To explore which EMG or FSR features explain behavioural performance, we ran exploratory backward stepwise regressions (entry: 0.05; removal: 0.1).

To see if participants were neutral or experienced positive/negative embodiment towards the Thumb, a one-sample t-test was used for each sub-group to see if the scores in each category were different from zero. An independent samples t-test was then used to compare the sub-groups.

## Acknowledgments

The study was funded by UKRI’s Frontier Research Guarantee (EP/X040372/1), and the Medical Research Council (MC_uu_00030/10). J.P.R. was funded by the Engineering and Physical Sciences Research Council studentship. T.R.M. was partially funded by a Wellcome Trust Senior Research Fellowship (215575/Z/19/Z). We acknowledge support from Meta through a charitable gift. We thank our participants for their valuable help. We thank Clara Gallay for data collection. We thank Mario Bracklein (CTRL-labs at Reality Labs) for helpful insight on the manuscript.

For the purpose of open access, the authors have applied a Creative Commons Attribution (CC BY) licence to any Author Accepted Manuscript version arising from this submission.

## Authors Contributions

J.P.R. and T.R.M. designed the study. J.P.R., F.C. and K.G. made the technical developments. J.P.R. collected and analysed the data. F.C., K.G. and M.Z. provided assistance with data collection. M.Z., helped with recruitment. J.P.R., L.D. and T.R.M. interpreted the results. J.P.R. wrote the manuscripts. L.D., H.D., D.C. and T.R.M. provided helpful comments for the manuscripts.

## References

Prattichizzo, D. et al.(2021) ‘Human augmentation by wearable supernumerary robotic limbs: review and perspectives’, Progress in Biomedical Engineering, 3(4), p. 042005. Available at: 10.1088/2516-1091/ac2294.

Eden, J. et al.(2022) ‘Principles of human movement augmentation and the challenges in making it a reality’, Nature Communications, 13(1), p. 1345. Available at: 10.1038/s41467-022-28725-7.

Dominijanni, G. et al.(2021) ‘The neural resource allocation problem when enhancing human bodies with extra robotic limbs’, Nature Machine Intelligence, 3(10), pp. 850–860. Available at: 10.1038/s42256-021-00398-9.

Mooney, L.M. and Herr, H.M. (2016) ‘Biomechanical walking mechanisms underlying the metabolic reduction caused by an autonomous exoskeleton’, Journal of NeuroEngineering and Rehabilitation, 13(1), p. 4. Available at: 10.1186/s12984-016-0111-3.

Kim, J. et al.(2019) ‘Reducing the metabolic rate of walking and running with a versatile, portable exosuit’, Science, 365(6454), pp. 668–672. Available at: 10.1126/science.aav7536.

Waight, J.L. et al.(2023) ‘From functional neuroimaging to neurostimulation: fNIRS devices as cognitive enhancers’, Behavior Research Methods, 56(3), pp. 2227–2242. Available at: 10.3758/s13428-023-02144-y.

Follmann, A. et al.(2019) ‘Technical Support by Smart Glasses During a Mass Casualty Incident: A Randomized Controlled Simulation Trial on Technically Assisted Triage and Telemedical App Use in Disaster Medicine’, Journal of Medical Internet Research, 21(1), p. e11939. Available at: 10.2196/11939.

Wu, F. and Asada, H. (2014) ‘Bio-Artificial Synergies for Grasp Posture Control of Supernumerary Robotic Fingers’, in Robotics: Science and Systems X. Robotics: Science and Systems 2014, Robotics: Science and Systems Foundation. Available at: 10.15607/RSS.2014.X.027.

Makin, T.R., Micera, S. and Miller, L.E. (2022) ‘Neurocognitive and motor-control challenges for the realization of bionic augmentation’, Nature Biomedical Engineering, 7(4), pp. 344–348. Available at: 10.1038/s41551-022-00930-1.

Verdel, D. et al.(2024) ‘A predictive coding framework for safe and versatile control of supernumerary robotic limbs’, hal-04830213 [Preprint].

Arozi, M. et al.(2020) ‘Pattern Recognition of Single-Channel sEMG Signal Using PCA and ANN Method to Classify Nine Hand Movements’, Symmetry, 12(4), p. 541. Available at: 10.3390/sym12040541.

Hahne, J.M. et al.(2018) ‘Simultaneous control of multiple functions of bionic hand prostheses: Performance and robustness in end users’, Science Robotics, 3(19), p. eaat3630. Available at: 10.1126/scirobotics.aat3630.

Hussain, I., Spagnoletti, G., et al. (2016) ‘An EMG Interface for the Control of Motion and Compliance of a Supernumerary Robotic Finger’, Frontiers in Neurorobotics, 10. Available at: 10.3389/fnbot.2016.00018.

Meraz, N.S., Shikida, H. and Hasegawa, Y. (2017) ‘Auricularis muscles based control interface for robotic extra thumb’, in 2017 International Symposium on Micro-NanoMechatronics and Human Science (MHS). 2017 International Symposium on Micro-NanoMechatronics and Human Science (MHS), Nagoya: IEEE, pp. 1–3. Available at: 10.1109/MHS.2017.8305192.

Tavakoli, M., Benussi, C. and Lourenco, J.L. (2017) ‘Single channel surface EMG control of advanced prosthetic hands: A simple, low cost and efficient approach’, Expert Systems with Applications, 79, pp. 322–332. Available at: 10.1016/j.eswa.2017.03.012.

Yang, D. et al.(2014) ‘Experimental Study of an EMG-Controlled 5-DOF Anthropomorphic Prosthetic Hand for Motion Restoration’, Journal of Intelligent & Robotic Systems, 76(3–4), pp. 427–441. Available at: 10.1007/s10846-014-0037-6.

Trigili, E. et al.(2019) ‘Detection of movement onset using EMG signals for upper-limb exoskeletons in reaching tasks’, Journal of NeuroEngineering and Rehabilitation, 16(1), p. 45. Available at: 10.1186/s12984-019-0512-1.

Carvalho, C.R. et al.(2023) ‘Review of electromyography onset detection methods for real-time control of robotic exoskeletons’, Journal of NeuroEngineering and Rehabilitation, 20(1), p. 141. Available at: 10.1186/s12984-023-01268-8.

Kiguchi, K. and Hayashi, Y. (2012) ‘An EMG-Based Control for an Upper-Limb Power-Assist Exoskeleton Robot’, IEEE Transactions on Systems, Man, and Cybernetics, Part B (Cybernetics), 42(4), pp. 1064–1071. Available at: 10.1109/TSMCB.2012.2185843.

CTRL-labs at Reality Labs et al. (2024) ‘A generic noninvasive neuromotor interface for human-computer interaction’. Available at: 10.1101/2024.02.23.581779.

Jie, J. et al.(2021) ‘High dimensional feature data reduction of multichannel sEMG for gesture recognition based on double phases PSO’, Complex & Intelligent Systems, 7(4), pp. 1877–1893. Available at: 10.1007/s40747-020-00232-6.

Kieliba, P. et al.(2021) ‘Robotic hand augmentation drives changes in neural body representation’, Science Robotics, 6(54), p. eabd7935. Available at: 10.1126/scirobotics.abd7935.

Bräcklein, M. et al.(2022) ‘The control and training of single motor units in isometric tasks are constrained by a common input signal’, eLife, 11, p. e72871. Available at: 10.7554/eLife.72871.

Formento, E., Botros, P. and Carmena, J.M. (2021) ‘Skilled independent control of individual motor units via a non-invasive neuromuscular–machine interface’, Journal of Neural Engineering, 18(6), p. 066019. Available at: 10.1088/1741-2552/ac35ac.

Clode, D. et al.(2024) ‘Evaluating initial usability of a hand augmentation device across a large and diverse sample’, Science Robotics, 9(90), p. eadk5183. Available at: 10.1126/scirobotics.adk5183.

Amoruso, E. et al.(2022) ‘Intrinsic somatosensory feedback supports motor control and learning to operate artificial body parts’, Journal of Neural Engineering, 19(1), p. 016006. Available at: 10.1088/1741-2552/ac47d9.

Schone, H.R. et al.(2024) ‘Biomimetic versus arbitrary motor control strategies for bionic hand skill learning’, Nature Human Behaviour, 8(6), pp. 1108–1123. Available at: 10.1038/s41562-023-01811-6.

Clemente, F. et al.(2016) ‘Non-Invasive, Temporally Discrete Feedback of Object Contact and Release Improves Grasp Control of Closed-Loop Myoelectric Transradial Prostheses’, IEEE Transactions on Neural Systems and Rehabilitation Engineering, 24(12), pp. 1314– 1322. Available at: 10.1109/TNSRE.2015.2500586.

Witteveen, H.J.B. et al.(2012) ‘Hand-opening feedback for myoelectric forearm prostheses: Performance in virtual grasping tasks influenced by different levels of distraction’, The Journal of Rehabilitation Research and Development, 49(10), p. 1517. Available at: 10.1682/JRRD.2011.12.0243.

Guthrie, M.D. et al.(2022) ‘The impact of distractions on intracortical brain–computer interface control of a robotic arm’, Brain-Computer Interfaces, 9(1), pp. 23–35. Available at: 10.1080/2326263X.2021.1980292.

Longo, M.R. et al.(2008) ‘What is embodiment? A psychometric approach’, Cognition, 107(3), pp. 978–998. Available at: 10.1016/j.cognition.2007.12.004.

Wetzels, R. and Wagenmakers, E.-J. (2012) ‘A default Bayesian hypothesis test for correlations and partial correlations’, Psychonomic Bulletin & Review, 19(6), pp. 1057–1064. Available at: 10.3758/s13423-012-0295-x.

Makin, T.R. and Orban De Xivry, J.-J. (2019) ‘Ten common statistical mistakes to watch out for when writing or reviewing a manuscript’, eLife, 8, p. e48175. Available at: 10.7554/eLife.48175.

Scott, S.H. et al.(2015) ‘Feedback control during voluntary motor actions’, Current Opinion in Neurobiology, 33, pp. 85–94. Available at: 10.1016/j.conb.2015.03.006.

d’Avella, A., Saltiel, P. and Bizzi, E. (2003) ‘Combinations of muscle synergies in the construction of a natural motor behavior’, Nature Neuroscience, 6(3), pp. 300–308. Available at: 10.1038/nn1010.

Farina, D., Fosci, M. and Merletti, R. (2002) ‘Motor unit recruitment strategies investigated by surface EMG variables’, Journal of Applied Physiology, 92(1), pp. 235–247. Available at: 10.1152/jappl.2002.92.1.235.

Stegeman, D.F. et al.(2000) ‘Surface EMG models: properties and applications’, Journal of Electromyography and Kinesiology, 10(5), pp. 313–326. Available at: 10.1016/S1050-6411(00)00023-7.

Wong, Y.-M. and Ng, G.Y.F. (2006) ‘Surface electrode placement affects the EMG recordings of the quadriceps muscles’, Physical Therapy in Sport, 7(3), pp. 122–127. Available at: 10.1016/j.ptsp.2006.03.006.

Raviv, L., Lupyan, G. and Green, S.C. (2022) ‘How variability shapes learning and generalization’, Trends in Cognitive Sciences, 26(6), pp. 462–483. Available at: 10.1016/j.tics.2022.03.007.

