## Supplementary_materials for "Muscle control of an extra robotic digit"

| Variable | | FS mean | FS standard deviation | FS variability | FS fractal dimension | FS 1st derivative variance | FS 2nd derivative variance | FS slope change | FS waveform length |
|---|---|---|---|---|---|---|---|---|---|
| 1. FS mean | Pearson's r | — | | | | | | | |
| | p-value | — | | | | | | | |
| 2. FS standard deviation | Pearson's r | −0.733 | — | | | | | | |
| | p-value | < .001 | — | | | | | | |
| 3. FS variability | Pearson's r | −0.921 | 0.856 | — | | | | | |
| | p-value | < .001 | < .001 | — | | | | | |
| 4. FS fractal dimension | Pearson's r | −0.268 | 0.459 | 0.337 | — | | | | |
| | p-value | 0.267 | 0.048 | 0.158 | — | | | | |
| 5. FS 1st derivative variance | Pearson's r | −0.569 | 0.674 | 0.645 | −0.135 | — | | | |
| | p-value | 0.011 | 0.002 | 0.003 | 0.583 | — | | | |
| 6. FS 2nd derivative variance | Pearson's r | −0.403 | 0.545 | 0.468 | −0.271 | 0.975 | — | | |
| | p-value | 0.087 | 0.016 | 0.043 | 0.262 | < .001 | — | | |
| 7. FS slope change | Pearson's r | 0.412 | −0.033 | −0.357 | −0.197 | 0.052 | 0.150 | — | |
| | p-value | 0.080 | 0.895 | 0.134 | 0.418 | 0.832 | 0.540 | — | |
| 8. FS waveform length | Pearson's r | −0.530 | 0.675 | 0.578 | −0.176 | 0.859 | 0.844 | 0.337 | — |
| | p-value | 0.020 | 0.002 | 0.010 | 0.470 | < .001 | < .001 | 0.158 | — |

**Table S1.** Correlation (Pearson's) table between the eight FS control signal parameter, extracted during the post-training Proportional Control task. Parameters with similar profiles (determined based on high correlation with other parameters of interest) were disregarded to minimise covariance.

| Variable | | EMG mean | EMG standard deviation | EMG variability | EMG fractal dimension | EMG 1st derivative variance | EMG 2nd derivative variance | EMG slope change | EMG waveform length |
|---|---|---|---|---|---|---|---|---|---|
| 1. EMG mean | Pearson's r | — | | | | | | | |
| | p-value | — | | | | | | | |
| 2. EMG standard deviation | Pearson's r | 0.584 | — | | | | | | |
| | p-value | 0.003 | — | | | | | | |
| 3. EMG variability | Pearson's r | −0.490 | 0.268 | — | | | | | |
| | p-value | 0.015 | 0.205 | — | | | | | |
| 4. EMG fractal dimension | Pearson's r | −0.033 | 0.363 | 0.456 | — | | | | |
| | p-value | 0.880 | 0.081 | 0.025 | — | | | | |
| 5. EMG 1st derivative variance | Pearson's r | 0.783 | 0.871 | −0.023 | 0.117 | — | | | |
| | p-value | < .001 | < .001 | 0.916 | 0.587 | — | | | |
| 6. EMG 2nd derivative variance | Pearson's r | 0.724 | 0.810 | −0.056 | −0.015 | 0.971 | — | | |
| | p-value | < .001 | < .001 | 0.796 | 0.946 | < .001 | — | | |
| 7. EMG slope change | Pearson's r | 0.771 | 0.825 | −0.075 | 0.102 | 0.838 | 0.823 | — | |
| | p-value | < .001 | < .001 | 0.727 | 0.636 | < .001 | < .001 | — | |
| 8. EMG waveform length | Pearson's r | 0.719 | 0.917 | 0.046 | 0.173 | 0.948 | 0.935 | 0.941 | — |
| | p-value | < .001 | < .001 | 0.833 | 0.420 | < .001 | < .001 | < .001 | — |

**Table S2.** Correlation (Pearson's) table between the eight EMG control signal parameter, extracted during the post-training Proportional Control task. Parameters with similar profiles (determined based on high correlation with other parameters of interest) were disregarded to minimise covariance.

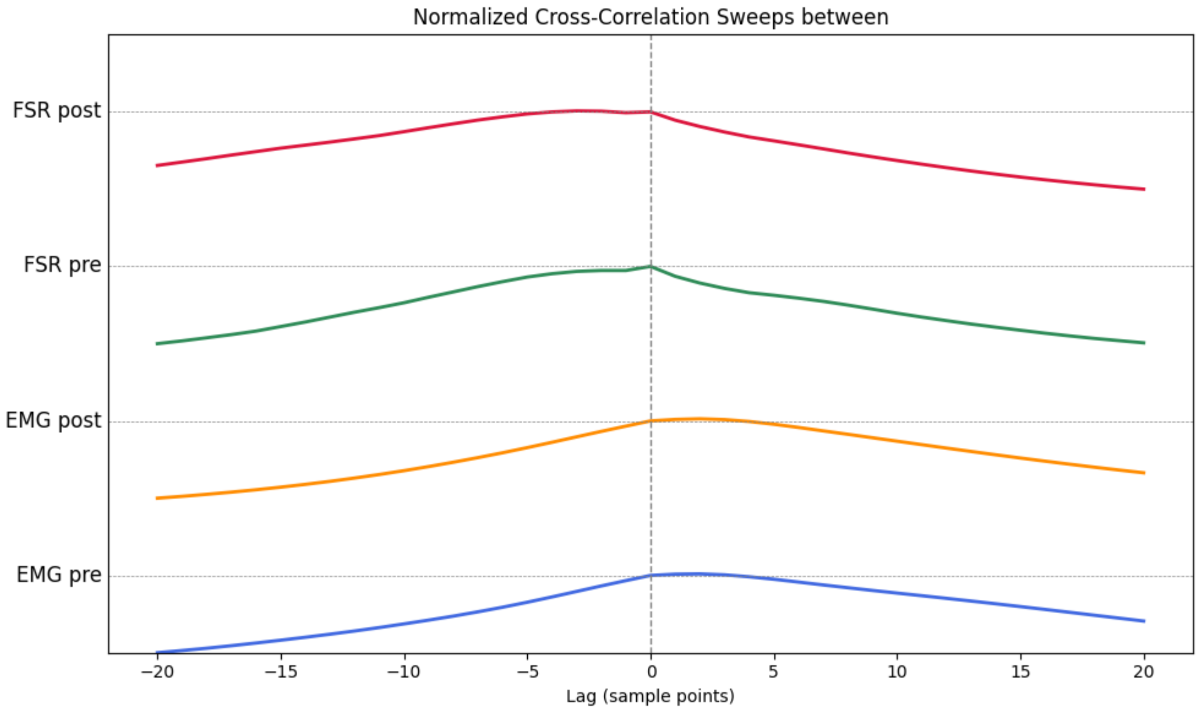

**Figure S1.** Normalized cross-correlation sweeps. For data extracted during the FS Proportional Control task (pre: green; post: red), the higher correlation values to the left of the zero suggests the EMG amplitude fluctuations slightly precede the FS signal, but the lack of a clear peak outside the zero line implies this lag may not be consistent across participants. For EMG pre and post (blue and orange), the positive peak to the right supports the FS signal occurs slightly later (about 2 sample point) than the EMG signal. Beyond a demonstration a temporal advantage to the EMG signals, which consistently precede the FS signal, these findings indicate that during EMG control, participants tend to press the toes even when it's not required for FS input, thus strengthens the motor mapping that benefits later FS control.

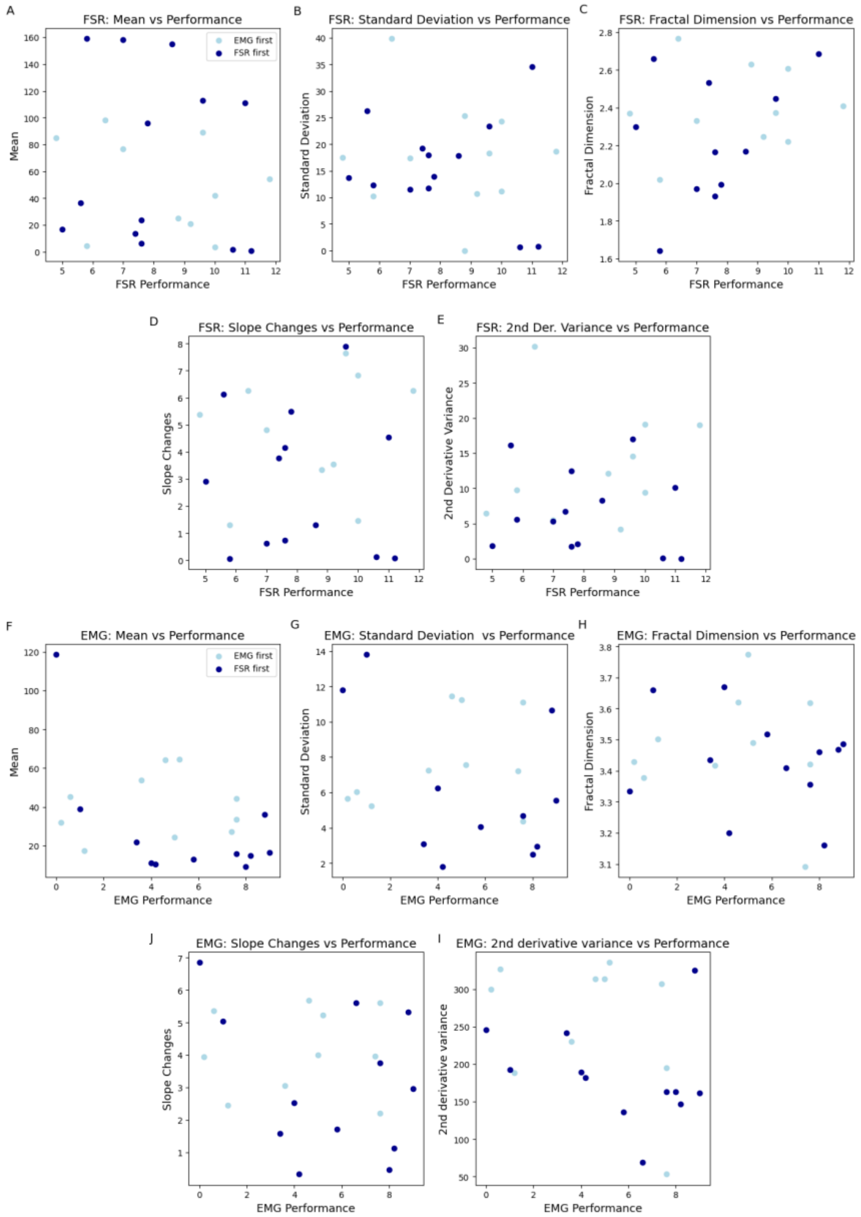


**Figure S2. EMG and FS parameters used for predictive modelling against performance.** Panels show the five signal parameters extracted from force (FS; A-E) and EMG (F-J) control signals, extracted from the post-training Proportional Control task. These features are plotted against motor performance using the related control modality. (A, F) mean signal amplitude. (B, G) signal's standard deviation. (C, H) fractal dimension. (D, I) slope change (J, E) second derivative variance. These features capture different aspects of signal stability, variability, and temporal dynamics, and were used in regression models to predict task performance.

Model Summary – FSR behavioural

| Model | R | $R^2$ | Adjusted $R^2$ | RMSE |
|---|---|---|---|---|
| 1 | 0.843 | 0.710 | 0.526 | 1.350 |
| 2 | 0.842 | 0.710 | 0.565 | 1.294 |
| 3 | 0.836 | 0.699 | 0.583 | 1.267 |

ANOVA ▼

| Model | | Sum of Squares | df | Mean Square | F | p |
|---|---|---|---|---|---|---|
| 1 | Regression | 49.143 | 7 | 7.020 | 3.852 | 0.023 |
| | Residual | 20.048 | 11 | 1.823 | | |
| | Total | 69.192 | 18 | | | |
| 2 | Regression | 49.110 | 6 | 8.185 | 4.891 | 0.009 |
| | Residual | 20.082 | 12 | 1.673 | | |
| | Total | 69.192 | 18 | | | |
| 3 | Regression | 48.335 | 5 | 9.667 | 6.025 | 0.004 |
| | Residual | 20.857 | 13 | 1.604 | | |
| | Total | 69.192 | 18 | | | |

Coefficients ▼

| Model | | Unstandardized | Standard Error | Standardized | t | p |
|---|---|---|---|---|---|---|
| 1 | (Intercept) | 50.923 | 55.717 | | 0.914 | 0.380 |
| | FS mean | −0.009 | 0.010 | −0.319 | −0.922 | 0.376 |
| | FS standard deviation | −0.004 | 0.027 | −0.060 | −0.135 | 0.895 |
| | FS slope change | −3.136 | 5.836 | −0.138 | −0.537 | 0.602 |
| | FS fractal dimension | −4.260 | 2.575 | −0.462 | −1.655 | 0.126 |
| | FS 2nd derivative variance | −0.004 | 0.002 | −0.622 | −1.971 | 0.074 |
| | EMG behavioural | 0.353 | 0.144 | 0.504 | 2.451 | 0.032 |
| | control order | −2.379 | 0.824 | −0.579 | −2.886 | 0.015 |
| 2 | (Intercept) | 53.991 | 48.744 | | 1.108 | 0.290 |
| | FS mean | −0.008 | 0.007 | −0.285 | −1.249 | 0.235 |
| | FS slope change | −3.463 | 5.088 | −0.152 | −0.681 | 0.509 |
| | FS fractal dimension | −4.518 | 1.654 | −0.490 | −2.731 | 0.018 |
| | FS 2nd derivative variance | −0.004 | 0.001 | −0.652 | −2.986 | 0.011 |
| | EMG behavioural | 0.362 | 0.125 | 0.516 | 2.898 | 0.013 |
| | control order | −2.404 | 0.769 | −0.586 | −3.127 | 0.009 |
| 3 | (Intercept) | 20.936 | 4.136 | | 5.062 | < .001 |
| | FS mean | −0.011 | 0.006 | −0.370 | −1.976 | 0.070 |
| | FS fractal dimension | −4.609 | 1.614 | −0.500 | −2.855 | 0.014 |
| | FS 2nd derivative variance | −0.005 | 0.001 | −0.715 | −3.690 | 0.003 |
| | EMG behavioural | 0.371 | 0.121 | 0.529 | 3.058 | 0.009 |
| | control order | −2.120 | 0.632 | −0.516 | −3.354 | 0.005 |

**Table S3.** Regression output for the backward stepwise regression with FS signal and EMG behavioural parameters as a predictor of force control performance.

Model Summary – EMG behavioural

| Model | R | $R^2$ | Adjusted $R^2$ | RMSE |
|---|---|---|---|---|
| 1 | 0.492 | 0.242 | −0.090 | 3.013 |
| 2 | 0.492 | 0.242 | −0.026 | 2.923 |
| 3 | 0.492 | 0.242 | 0.031 | 2.841 |
| 4 | 0.491 | 0.241 | 0.081 | 2.766 |
| 5 | 0.435 | 0.189 | 0.068 | 2.786 |
| 6 | 0.421 | 0.177 | 0.099 | 2.740 |
| 7 | 0.339 | 0.115 | 0.075 | 2.776 |
| 8 | 0.000 | 0.000 | 0.000 | 2.886 |

ANOVA

| Model | | Sum of Squares | df | Mean Square | F | p |
|---|---|---|---|---|---|---|
| 1 | Regression | 46.313 | 7 | 6.616 | 0.729 | 0.651 |
| | Residual | 145.245 | 16 | 9.078 | | |
| | Total | 191.558 | 23 | | | |
| 2 | Regression | 46.301 | 6 | 7.717 | 0.903 | 0.515 |
| | Residual | 145.257 | 17 | 8.545 | | |
| | Total | 191.558 | 23 | | | |
| 3 | Regression | 46.290 | 5 | 9.258 | 1.147 | 0.372 |
| | Residual | 145.269 | 18 | 8.070 | | |
| | Total | 191.558 | 23 | | | |
| 4 | Regression | 46.198 | 4 | 11.550 | 1.510 | 0.239 |
| | Residual | 145.360 | 19 | 7.651 | | |
| | Total | 191.558 | 23 | | | |
| 5 | Regression | 36.272 | 3 | 12.091 | 1.557 | 0.231 |
| | Residual | 155.287 | 20 | 7.764 | | |
| | Total | 191.558 | 23 | | | |
| 6 | Regression | 33.927 | 2 | 16.963 | 2.260 | 0.129 |
| | Residual | 157.632 | 21 | 7.506 | | |
| | Total | 191.558 | 23 | | | |
| 7 | Regression | 21.988 | 1 | 21.988 | 2.853 | 0.105 |
| | Residual | 169.570 | 22 | 7.708 | | |
| | Total | 191.558 | 23 | | | |

Coefficients ▼

| Model | | Unstandardized | Standard Error | Standardized | t | p |
|---|---|---|---|---|---|---|
| 1 | (Intercept) | −0.486 | 19.009 | | −0.026 | 0.980 |
| | EMG mean | −0.041 | 0.043 | −0.352 | −0.944 | 0.359 |
| | EMG standard deviation | 0.052 | 0.512 | 0.065 | 0.101 | 0.921 |
| | EMG fractal dimension | −0.260 | 5.454 | −0.014 | −0.048 | 0.963 |
| | EMG slope change | 0.530 | 0.834 | 0.353 | 0.635 | 0.534 |
| | EMG 2nd derivative variance | −0.122 | 3.366 | −0.022 | −0.036 | 0.972 |
| | control order | 1.626 | 1.487 | 0.287 | 1.093 | 0.290 |
| | FS behavioural | 0.556 | 0.374 | 0.393 | 1.487 | 0.156 |
| 2 | (Intercept) | −0.694 | 17.584 | | −0.039 | 0.969 |
| | EMG mean | −0.041 | 0.039 | −0.356 | −1.053 | 0.307 |
| | EMG standard deviation | 0.039 | 0.361 | 0.049 | 0.108 | 0.915 |
| | EMG fractal dimension | −0.177 | 4.795 | −0.010 | −0.037 | 0.971 |
| | EMG slope change | 0.529 | 0.809 | 0.352 | 0.654 | 0.522 |
| | control order | 1.641 | 1.380 | 0.289 | 1.189 | 0.251 |
| | FS behavioural | 0.550 | 0.323 | 0.388 | 1.701 | 0.107 |
| 3 | (Intercept) | −1.332 | 3.052 | | −0.436 | 0.668 |
| | EMG mean | −0.041 | 0.038 | −0.355 | −1.085 | 0.292 |
| | EMG standard deviation | 0.032 | 0.304 | 0.041 | 0.106 | 0.916 |
| | EMG slope change | 0.538 | 0.753 | 0.358 | 0.714 | 0.484 |
| | control order | 1.652 | 1.313 | 0.291 | 1.258 | 0.225 |
| | FS behavioural | 0.554 | 0.296 | 0.391 | 1.871 | 0.078 |
| 4 | (Intercept) | −1.281 | 2.935 | | −0.437 | 0.667 |
| | EMG mean | −0.042 | 0.036 | −0.360 | −1.139 | 0.269 |
| | EMG slope change | 0.598 | 0.491 | 0.398 | 1.218 | 0.238 |
| | control order | 1.687 | 1.238 | 0.297 | 1.363 | 0.189 |
| | FS behavioural | 0.550 | 0.286 | 0.388 | 1.923 | 0.070 |
| 5 | (Intercept) | −0.947 | 2.942 | | −0.322 | 0.751 |
| | EMG slope change | 0.181 | 0.329 | 0.120 | 0.550 | 0.589 |
| | control order | 1.691 | 1.247 | 0.298 | 1.356 | 0.190 |
| | FS behavioural | 0.513 | 0.286 | 0.362 | 1.793 | 0.088 |
| 6 | (Intercept) | −0.131 | 2.497 | | −0.052 | 0.959 |
| | control order | 1.420 | 1.126 | 0.250 | 1.261 | 0.221 |
| | FS behavioural | 0.508 | 0.281 | 0.359 | 1.807 | 0.085 |
| 7 | (Intercept) | 0.871 | 2.399 | | 0.363 | 0.720 |
| | FS behavioural | 0.480 | 0.284 | 0.339 | 1.689 | 0.105 |
| 8 | (Intercept) | 4.808 | 0.589 | | 8.162 | < .001 |

**Table S4.** Regression output for the backward stepwise regression with EMG signal and force control behavioural parameters as a predictor of muscle control performance.

Model Summary – EMG behavioural

| Model | R | $R^2$ | Adjusted $R^2$ | RMSE |
|---|---|---|---|---|
| 1 | 0.783 | 0.614 | 0.406 | 2.315 |
| 2 | 0.767 | 0.589 | 0.412 | 2.303 |
| 3 | 0.730 | 0.533 | 0.377 | 2.370 |
| 4 | 0.699 | 0.488 | 0.361 | 2.401 |

ANOVA

| Model | | Sum of Squares | df | Mean Square | F | p |
|---|---|---|---|---|---|---|
| 1 | Regression | 110.671 | 7 | 15.810 | 2.949 | 0.044 |
| | Residual | 69.699 | 13 | 5.361 | | |
| | Total | 180.370 | 20 | | | |
| 2 | Regression | 106.148 | 6 | 17.691 | 3.337 | 0.030 |
| | Residual | 74.221 | 14 | 5.302 | | |
| | Total | 180.370 | 20 | | | |
| 3 | Regression | 96.114 | 5 | 19.223 | 3.422 | 0.029 |
| | Residual | 84.256 | 15 | 5.617 | | |
| | Total | 180.370 | 20 | | | |
| 4 | Regression | 88.105 | 4 | 22.026 | 3.820 | 0.023 |
| | Residual | 92.265 | 16 | 5.767 | | |
| | Total | 180.370 | 20 | | | |

Coefficients

| Model | | Unstandardized | Standard Error | Standardized | t | p |
|---|---|---|---|---|---|---|
| 1 | (Intercept) | −653.743 | 227.974 | | −2.868 | 0.013 |
| | control order | 5.147 | 2.034 | 0.877 | 2.531 | 0.025 |
| | FS behavioural | 0.919 | 0.296 | 0.655 | 3.101 | 0.008 |
| | raw EMG fractal dimension | 4.030 | 4.388 | 0.299 | 0.918 | 0.375 |
| | raw EMG zero crossing | −0.066 | 0.023 | −1.723 | −2.899 | 0.012 |
| | raw EMG slope change | 0.348 | 0.119 | 2.511 | 2.924 | 0.012 |
| | raw EMG standard deviation | −0.111 | 0.061 | −1.234 | −1.839 | 0.089 |
| | raw EMG mean | −1.006 | 0.613 | −0.549 | −1.640 | 0.125 |
| 2 | (Intercept) | −511.649 | 166.497 | | −3.073 | 0.008 |
| | control order | 4.836 | 1.994 | 0.824 | 2.425 | 0.029 |
| | FS behavioural | 0.882 | 0.292 | 0.629 | 3.020 | 0.009 |
| | raw EMG zero crossing | −0.054 | 0.018 | −1.404 | −2.928 | 0.011 |
| | raw EMG slope change | 0.274 | 0.087 | 1.979 | 3.140 | 0.007 |
| | raw EMG standard deviation | −0.067 | 0.037 | −0.745 | −1.836 | 0.088 |
| | raw EMG mean | −0.739 | 0.537 | −0.403 | −1.376 | 0.191 |
| 3 | (Intercept) | −502.661 | 171.248 | | −2.935 | 0.010 |
| | control order | 5.552 | 1.982 | 0.946 | 2.802 | 0.013 |
| | FS behavioural | 0.971 | 0.293 | 0.693 | 3.314 | 0.005 |
| | raw EMG zero crossing | −0.052 | 0.019 | −1.365 | −2.771 | 0.014 |
| | raw EMG slope change | 0.262 | 0.089 | 1.892 | 2.932 | 0.010 |
| | raw EMG standard deviation | −0.033 | 0.028 | −0.367 | −1.194 | 0.251 |
| 4 | (Intercept) | −373.959 | 134.833 | | −2.773 | 0.014 |
| | control order | 5.448 | 2.006 | 0.928 | 2.716 | 0.015 |
| | FS behavioural | 0.885 | 0.288 | 0.632 | 3.076 | 0.007 |
| | raw EMG zero crossing | −0.036 | 0.013 | −0.926 | −2.789 | 0.013 |
| | raw EMG slope change | 0.194 | 0.070 | 1.398 | 2.787 | 0.013 |

**Table S5.** Regression output for the backward stepwise regression with raw EMG signal and force control behavioural parameters as a predictor of EMG performance.

Model Summary – FSR behavioural

| Model | R | $R^2$ | Adjusted $R^2$ | RMSE |
|---|---|---|---|---|
| 1 | 0.713 | 0.508 | 0.195 | 2.015 |
| 2 | 0.713 | 0.508 | 0.262 | 1.929 |
| 3 | 0.712 | 0.507 | 0.317 | 1.856 |
| 4 | 0.709 | 0.503 | 0.361 | 1.795 |
| 5 | 0.695 | 0.483 | 0.380 | 1.768 |

ANOVA

| Model | | Sum of Squares | df | Mean Square | F | p |
|---|---|---|---|---|---|---|
| 1 | Regression | 46.100 | 7 | 6.586 | 1.622 | 0.227 |
| | Residual | 44.667 | 11 | 4.061 | | |
| | Total | 90.766 | 18 | | | |
| 2 | Regression | 46.097 | 6 | 7.683 | 2.064 | 0.134 |
| | Residual | 44.670 | 12 | 3.722 | | |
| | Total | 90.766 | 18 | | | |
| 3 | Regression | 46.008 | 5 | 9.202 | 2.673 | 0.071 |
| | Residual | 44.758 | 13 | 3.443 | | |
| | Total | 90.766 | 18 | | | |
| 4 | Regression | 45.671 | 4 | 11.418 | 3.545 | 0.034 |
| | Residual | 45.096 | 14 | 3.221 | | |
| | Total | 90.766 | 18 | | | |
| 5 | Regression | 43.868 | 3 | 14.623 | 4.677 | 0.017 |
| | Residual | 46.898 | 15 | 3.127 | | |
| | Total | 90.766 | 18 | | | |

Coefficients

| Model | | Unstandardized | Standard Error | Standardized | t | p |
|---|---|---|---|---|---|---|
| 1 | (Intercept) | 206.761 | 152.932 | | 1.352 | 0.204 |
| | EMG behavioural | 0.532 | 0.198 | 0.660 | 2.686 | 0.021 |
| | control order | −3.813 | 2.016 | −0.871 | −1.892 | 0.085 |
| | raw EMG standard deviation | 0.002 | 0.014 | 0.071 | 0.150 | 0.883 |
| | raw EMG fractal dimension | −0.624 | 1.959 | −0.118 | −0.318 | 0.756 |
| | raw EMG mean | 0.032 | 1.154 | 0.006 | 0.028 | 0.978 |
| | raw EMG slope change | −0.103 | 0.079 | −1.060 | −1.306 | 0.218 |
| | raw EMG zero crossing | 0.007 | 0.011 | 0.320 | 0.644 | 0.533 |
| 2 | (Intercept) | 207.240 | 145.500 | | 1.424 | 0.180 |
| | EMG behavioural | 0.533 | 0.189 | 0.661 | 2.818 | 0.016 |
| | control order | −3.812 | 1.930 | −0.871 | −1.976 | 0.072 |
| | raw EMG standard deviation | 0.002 | 0.013 | 0.068 | 0.154 | 0.880 |
| | raw EMG fractal dimension | −0.622 | 1.874 | −0.117 | −0.332 | 0.746 |
| | raw EMG slope change | −0.103 | 0.075 | −1.060 | −1.364 | 0.198 |
| | raw EMG zero crossing | 0.007 | 0.010 | 0.321 | 0.677 | 0.511 |
| 3 | (Intercept) | 193.249 | 109.498 | | 1.765 | 0.101 |
| | EMG behavioural | 0.537 | 0.180 | 0.665 | 2.976 | 0.011 |
| | control order | −3.659 | 1.593 | −0.836 | −2.298 | 0.039 |
| | raw EMG fractal dimension | −0.442 | 1.414 | −0.083 | −0.313 | 0.759 |
| | raw EMG slope change | −0.096 | 0.057 | −0.987 | −1.670 | 0.119 |
| | raw EMG zero crossing | 0.006 | 0.009 | 0.288 | 0.707 | 0.492 |
| 4 | (Intercept) | 205.939 | 98.384 | | 2.093 | 0.055 |
| | EMG behavioural | 0.531 | 0.174 | 0.659 | 3.059 | 0.008 |
| | control order | −3.727 | 1.526 | −0.851 | −2.442 | 0.028 |
| | raw EMG slope change | −0.103 | 0.051 | −1.058 | −2.003 | 0.065 |
| | raw EMG zero crossing | 0.006 | 0.008 | 0.295 | 0.748 | 0.467 |
| 5 | (Intercept) | 142.721 | 49.613 | | 2.877 | 0.012 |
| | EMG behavioural | 0.482 | 0.158 | 0.597 | 3.047 | 0.008 |
| | control order | −3.019 | 1.180 | −0.690 | −2.558 | 0.022 |
| | raw EMG slope change | −0.070 | 0.025 | −0.716 | −2.743 | 0.015 |

**Table S6.** Regression output for the backward stepwise regression with raw EMG signal and force control behavioural parameters as a predictor of FS performance.

One Sample T-Test ▼

| | t | df | p |
|---|---|---|---|
| emg_body_ownership | −2.186 | 7 | 0.065 |
| fs_body_ownership | −2.063 | 10 | 0.066 |
| emg_agency | 2.458 | 7 | 0.044 |
| fs_agency | 3.525 | 10 | 0.005 |
| emg_somatosentation | −0.615 | 7 | 0.558 |
| fs_somatosentation | −0.770 | 10 | 0.459 |
| emg_body_image | −0.129 | 7 | 0.901 |
| fs_body_image | −0.243 | 10 | 0.813 |

**Table S7.** Individual sub-group statistics for the four embodiment categories, individual statistics are in accordance with combined.
